# Tks5 interactome reveals ER-associated machinery translation in invadosomes

**DOI:** 10.1101/2024.07.02.601728

**Authors:** Léa Normand, Benjamin Bonnard, Margaux Sala, Sylvaine Di-Tommaso, Cyril Dourthe, Anne-Aurélie Raymond, Jean-William Dupuy, Luc Mercier, Jacky G. Goetz, Violaine Moreau, Elodie Henriet, Frédéric Saltel

## Abstract

The ability to progress and invade through the extracellular matrix is a characteristic shared by both normal and cancer cells through the formation of structures called invadosomes gathering invadopodia and podosomes. These invadosomes are plastic and dynamic structures that can adopt different organizations depending on the cell types and the environment such as rosettes, dots or linear invadosomes. In this study, we used the specific invadosome marker Tks5 (SH3PXD2A), to identify common features in these different organizations. Tks5 immunoprecipitation coupled with mass spectrometry analysis allowed us to identify common proteins in these different models. We identified elements of the translation machinery, in particular the EIF4B protein, but also endoplasmic reticulum (ER) proteins as part of the invadosome structure. Providing new data on invadosome molecular composition through Tks5 interactome, we identified that ER-associated translation machinery is recruited to invadosome and involved in their formation, persistence and function in all types of invadosomes.

**Summary:** Invadosomes are invasive F-actin structures exhibiting different organizations that degrade the extracellular matrix. The study uses their universal marker, Tks5, to provide new data about invadosome molecular composition and reveal the role of ER-associated translation machinery in invadosome formation and function.

## Introduction

The ability to progress and invade through the surrounding environment is a characteristic shared by both normal and tumor cells. Extracellular matrix (ECM) breaching such as basement membrane or fibrillar collagen rich matrices is needed in physiological conditions such as embryogenesis or wound healing but is also required at the early stages of the metastatic cascade. During metastasis, cells will invade their surrounding matrix thanks to invadosomes. Invadosomes are dynamic actin-based structures that allow cells to interact and remodel the ECM through the recruitment of metalloproteinases^1,2^. Invadosome formation has been shown to depend on different factors such as soluble factors like growth factors^3–5^ or the mechanical properties^6,7^ and composition of the ECM^8^. These structures, classified as podosomes and invadopodia, respectively in normal cells and tumor cells, can adopt different organizations such as rosettes, aggregates or dots^6^. These plastic and dynamic structures are able to sense and adapt to their environment composition. Laminin can promote the formation of dots^9,10^, while fibrillar type I collagen matrix leads to a re-organization along the fibers forming linear invadosomes^9,11^ demonstrating a microenvironment-induced remodeling. This morphological plasticity is associated with a variation of the molecular composition of the structures: podosomes present a ring of adhesive proteins which is absent in invadopodia or linear invadosomes^11^. While podosomes and invadopodia are integrin-dependent structures, linear invadosome formation specifically depends on the discoidin domain receptor 1 (DDR1)^9,12^. On the other hand, actin-binding proteins like cortactin, the Arp2/3 complex, the neuronal Wiskott-Aldrich syndrome protein (N-WASP) and the GTPase Cdc42 are shared by all invadosome structures, but are also present in other actin structures^13–15^. However, the Tks5 scaffolding protein appears as a specific and common marker to all types of invadosomes^15^. Indeed, this protein is enriched in invadosomes but not in focal adhesion, filopodia, membrane ruffle or lamellipodia^16^. Tks5, a Src substrate protein, is required for invadosome formation^16^. Tks5 was discovered by Lock et *al.* in 1998 and is part of the p47 family^17^. It contains two major domains, the PX one that allows anchoring to the membrane, and the SH3 domain that allows protein-protein interaction. In addition to its major role in invadosome formation, Tks5 is also involved in cell division in bladder cancer and in cell proliferation via its action during the G1 phase of the cell cycle^18,19^. Tks5 belongs to a family containing also the Tsk4 protein involved in ECM degradation and in podosomes formation, while Tks5 is involved in all invadosome formation^20^.

Even if several studies over the last few years have aimed at better defining molecular composition involved in invadosome formation^21,22^, the common or specific markers for each structures in regard to their plasticity need to be investigated.

The aim of this article was to identify new and common molecular components present in all types of invadosome forms, namely rosettes, dots and linear invadosomes. To do so, we performed Tks5 interactome in different invadosome models. We used two cellular models, a murine fibroblast cell line expressing a constitutive activated form of the oncogene Src, NIH3T3-Src and a human epidermoid carcinoma cancer cell line, A431, this in two ECM contexts to generate rosettes, dots and linear invadosomes.

We identified 88 proteins commonly enriched in Tks5 interactome from four different types of invadosomes. While several identified proteins were already known to be involved in the formation or function of invadosomes (CD44, cortactin, ADAM15 or MMP14), we highlighted 54 new proteins not described to be involved into invadosome formation and/or function. Among these proteins, a large proportion was related to translation machinery confirming data obtained in a previous study^21^. We identified the eukaryotic translation initiation factor 4B (EIF4B) into invadosomes. We confirmed that translation and EIF4B are involved in the matrix degradation activity. Furthermore, we showed that the presence of endoplasmic reticulum (ER) proteins is required for invadosome persistence. These findings provide a better understanding of the invadosome regulation through the ER-associated translation machinery.

## Results

### Invadosomes variability and plasticity

In order to compare the different invadosome structures, we chose two complementary cellular models. The Src-transformed mouse fibroblast NIH-3T3-Src cell line that present invadosomes as rosettes when cultured on a gelatin substrate and the A431 human epidermoid carcinoma cell line which form invadopodia as dots^23^ (Figure 1a-b). As expected, the fibrillar collagen matrix led to their reorganization as linear invadosomes with a co-localization of F-actin and Tks5 along the collagen fibers in both cell lines^23^ (Figure 1a-b). These elements demonstrate the variety of shapes and plasticity of invadosomes. Thus, Tks5 is a universal marker of rosettes, dots and linear invadosomes, and is the best candidate to investigate common invadosome features.

**Figure 1:**
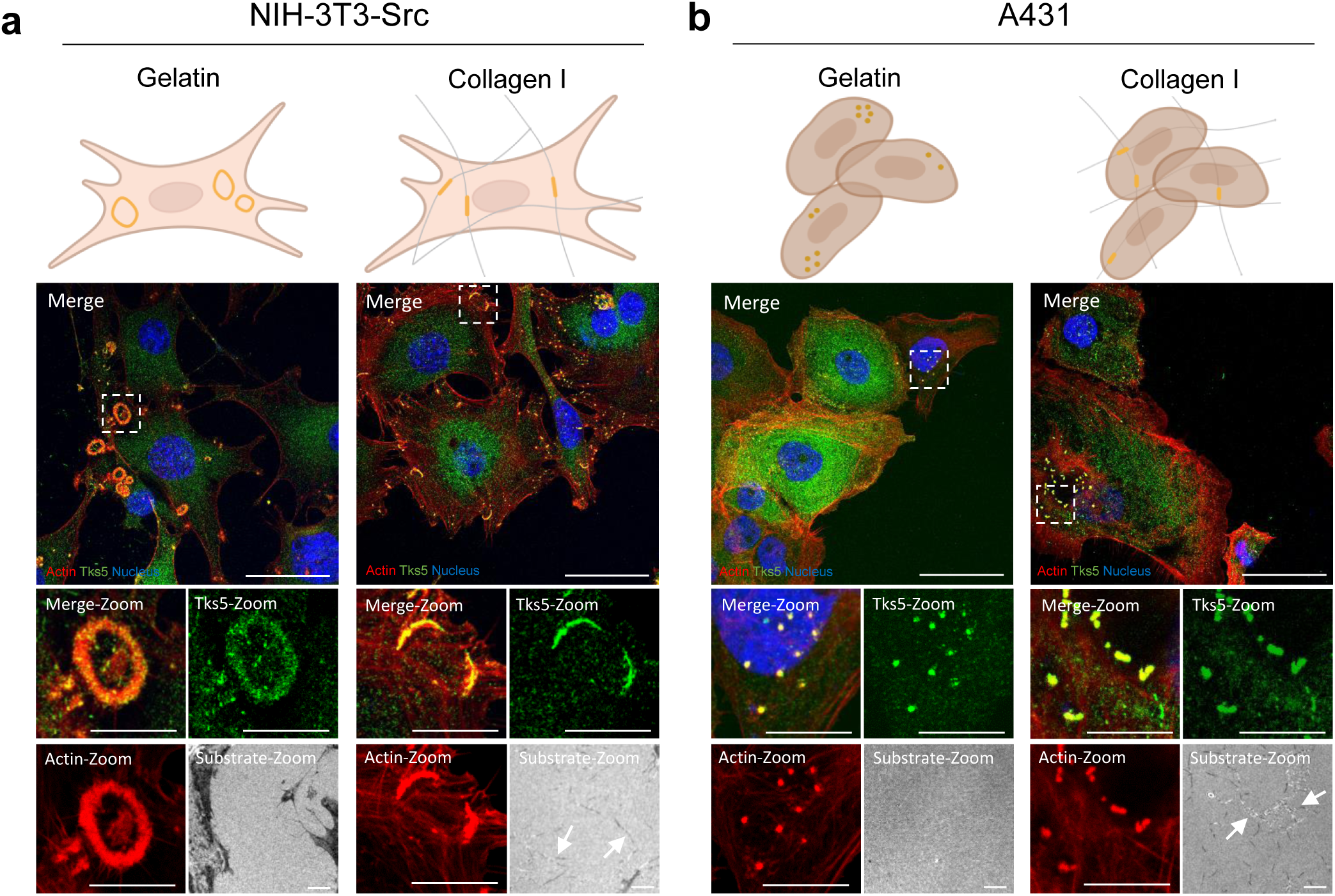
Invadosome plasticity. **a)** Confocal microscopy images of invadosome formation in NIH3T3-Src cells. The cells were seeded on gelatin or type I collagen to form rosettes and linear invadosomes respectively. Tks5 is stained in green, actin in red, nuclei in blue and collagen fibers were imaged by internal reflection microscopy (IRM). Scale bar: 40µm, zoom: 5µm. **b)** Confocal microscopy images of invadosome formation in A431 cells. The cells were seeded on gelatin or type I collagen to formed dots and linear invadosomes respectively. Tks5 is stained in green, actin in red, nuclei in blue and collagen was imaged by internal reflection microscopy (IRM). The arrows show the collagen fibers. Scale bar: 40µm, zoom: 5µm.

### Proteomics analysis of Tks5 interactome in invadosomes

To determine Tks5 interactome in these different invadosome types, we transfected NIH-3T3-Src and A431 cells with GFP-tagged Tks5 encoding plasmid. As expected, GFP-Tks5 colocalizes with endogenous Tks5 in all types of invadosomes (Figure 2a). We realized immunoprecipitation against GFP in both cell lines on plastic and type I collagen conditions (Supp Figure 1a). Indeed, on plastic, the cells behave as on a gelatin coating and thus form the same types of invadosomes, i.e. dots for A431 cells and rosettes for NIH-3T3-Src cells. Tks5 interactome was then assessed by LC-MS/MS mass spectrometry and the results are show in the Supplemental Table 1.

**Figure 2:**
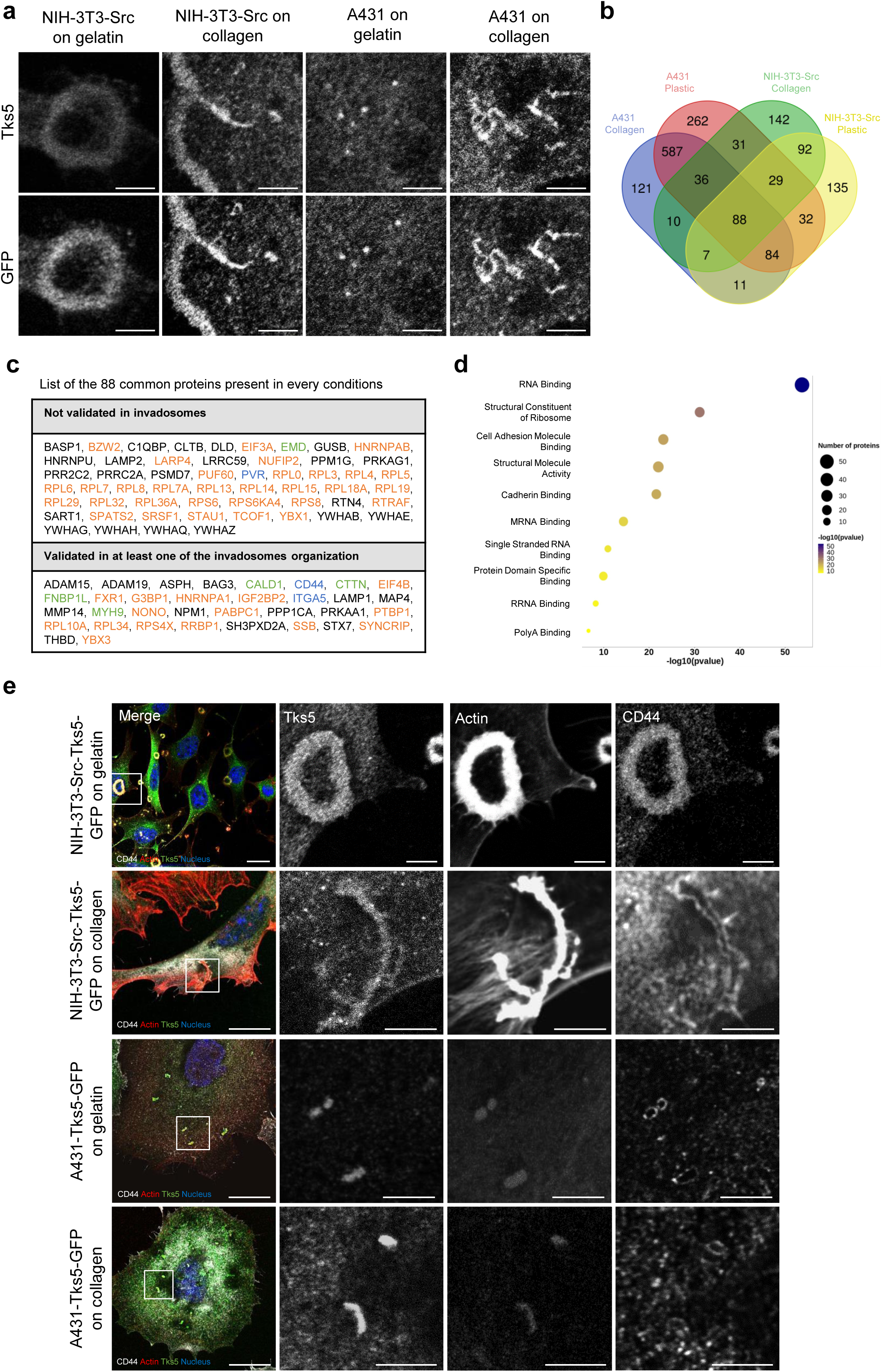
Proteomic analysis of Tks5 interactome in invadosomes. **a)** Confocal microscopy images of NIH3T3-Src-Tks5-GFP and A431-Tks5-GFP cells. The cells were seeded on gelatin and type I collagen to form invadosomes and stained for Tks5 and GFP. Scale bar: 5µm. **b)** Proteins interacting with Tks5-GFP by co-immunoprecipitation in NIH3T3-Src-Tks5-GFP and A431-Tks5-GFP cells on gelatin and type I collagen. 88 common proteins were identified in every conditions. **c)** List of the 88 commons proteins identified in every conditions. The color codes are in accordance with the table Supp Figure 2c: orange: translation, green: actin cytoskeleton and blue: adhesion. **d)** Bubble plot of the proteins related pathway. **e)** Confocal microscopy images of NIH-3T3-Src and A431-Tks5-GFP cells. The cells were seeded on gelatin or type I collagen and stained for Tks5 in green, actin in red, nuclei in blue and CD44 in grey. Scale bar: 20µm, zoom: 5µm.

We first determined the specific molecular signature associated with each invadosome organization (Supp Table 1-5). We commonly identified an enrichment in mitochondrial, endoplasmic reticulum (ER) and Golgi proteins (Supp Figure 1b-g). The gene set enrichment analysis (GSEA) performed on Tks5 interactome from linear invadosomes (A431 and NIH-3T3-Src cells) showed an enrichment for cell adhesion proteins such as CD34 and TSPAN6 absent in other invadosome organizations (Supp Figure 1f). These data show that depending on each organization, invadosomes exhibit specific proteins probably involved in their structure or function. Similarly, we also showed specific enrichment of translation proteins although each is organization specific: EIF5A in rosettes, RPS15 in dots or EIF2B4 in linear invadosomes (Supp Figure 1c, e, g).

We performed a comparative analysis with published data regarding proteins associated with Tks5 in 293T cells ^22^, and also identified cortactin (CTTN), the molecular chaperone Hsp90B1, the elongation factor EEF2 or the beta-actin protein (ACTB) (Supp Figure 2a, left panel). Similarly, another recent study that used proximity-labelling proteomic assay to identify Tks5 interactome in invadopodia in MDA-MB-231 cells, also identified CTTN, the microtubule-associated protein 4 (MAP4), the reticulon 4 protein (RTN4), the metalloproteinase ADAM15 and the eukaryotic translation initiation factor 4A3 (EIF4A3)^24^ (Supp Figure 2a, right panel). Since cortactin is well known to be present in all types of invadosomes, it can be used as a positive control to our approach^25,26^.

We then focused on proteins that were common to all invadosomes and identified 88 proteins present in dots, rosettes and linear invadosomes in the two cell types (Figure 2b-d). Among the 88 common proteins, 34 proteins such as CTTN, LAMP1, ADAM15 or MMP14 were previously identified in literature as components of in invadosomes, validating our mass spectrometry experiment. In order to validate identified proteins known to interact with Tks5^24,27,28^, we stained by immunofluorescence the receptor CD44 in all types of invadosomes and confirmed the co-localization of CD44 with Tks5 in all organizations of invadosomes (Figure 2e). Interestingly, the pattern of CD44 is different in rosettes where it colocalize with the actin, in comparison with dots and linear invadosome where CD44 surrounds the actin. Similarly, MAP4 co-localized with Tks5 in dots and linear invadosomes in A431 cells (Supp Figure 2b). These results were obtained from cells overexpressing Tks5-GFP but were also confirmed in WT cells, not overexpressing Tks5-GFP (data not shown). These experiments confirm the correct co-localization between Tks5 and the proteins CD44 and MAP4, identified in Tks5 interactome by mass spectrometry analysis.

However, a large proportion of identified protein has never been described to be present in invadosomes such as RPL19, RTN4 or EIF3A (Figure 2c). The GSEA performed on the 88 common proteins showed an enrichment in translation signaling (52% of total proteins) with an enrichment of RNA binding and structural constituent of ribosomes proteins (Figure 2d), including determinant elements of translation machinery such as RPL19, EIF4B or RPS6 (Supp Figure 2c). RPS6 is a component of the 40S subunit of the ribosome that allows the reading of mRNA^29^. EIF4B, on the other hand, is a translation factor that is part of the pre-initiation complex^30^. This complex, in association with the 40S subunit, allows to start mRNA translation. This result confirms the presence of the translation machinery at invadosomes, already identified by our team in NIH-3T3-Src cells using proteomics on laser captured rosette-like invadosomes (Ezzoukhry et *al.)*.

All together these results validate our experimental approach leading to the identification of new Tks5 partners into all types of invadosomes and suggest a link between Tks5 and the translation machinery.

### The translation machinery is present in all types of invadosomes

As we identified the translation machinery as a common feature shared by all the invadosome structures, we decided to focus our analysis on these translation proteins. Interestingly, other molecular studies of invadosomes, whether global or linked to Tks5, have identified proteins linked to translation^21,22,24,31^ (Supp Figure 2d). Briefly, Mallawaaratchy et *al.* identified proteins enriched in invadopodia of glioblastoma cells by mass spectrometry, Ezzoukhry et *al.* identified translation proteins enriched in invadosomes by laser microdissection combined with mass spectrometry, Stylli et *al.* identified proteins linked to Tks5 by co-immunoprecipitation and mass spectrometry in dots of 293T cells and Thuault et *al.* identified proteins linked to Tks5 by biotynilation technique in dots of MDA-MB-231. The RPS6 staining and the co-localization with Tks5 validated the presence of ribosomal proteins in all invadosomes (Figures 3a, 3b). We also showed the co-localization of the translational initiation factor, EIF4B, which was previously found enriched in rosettes^21^, with Tks5 in all organizations of invadosomes in both cell lines (Figure 3c and 3d). The presence of ribosomal proteins and translation factors in all invadosomes supports the hypothesis on an active translation in all these structures. Moreover, comparative analysis, using dataset obtained from Tks5 interactome from NIH-3T3-Src cells harboring rosettes and published rosettes proteome^21^, showed an enrichment in ribosomal and translation proteins (Supp Figure 2e-f). We commonly identified 79 proteins including 39 related to translation proteins (Supp Figure 2f). Among these 39 proteins, 20 are ribosomal proteins (Supp Figure 2f) which confirms the presence of ribosomes into invadosome rosettes as observed in Ezzoukhry et al.

**Figure 3:**
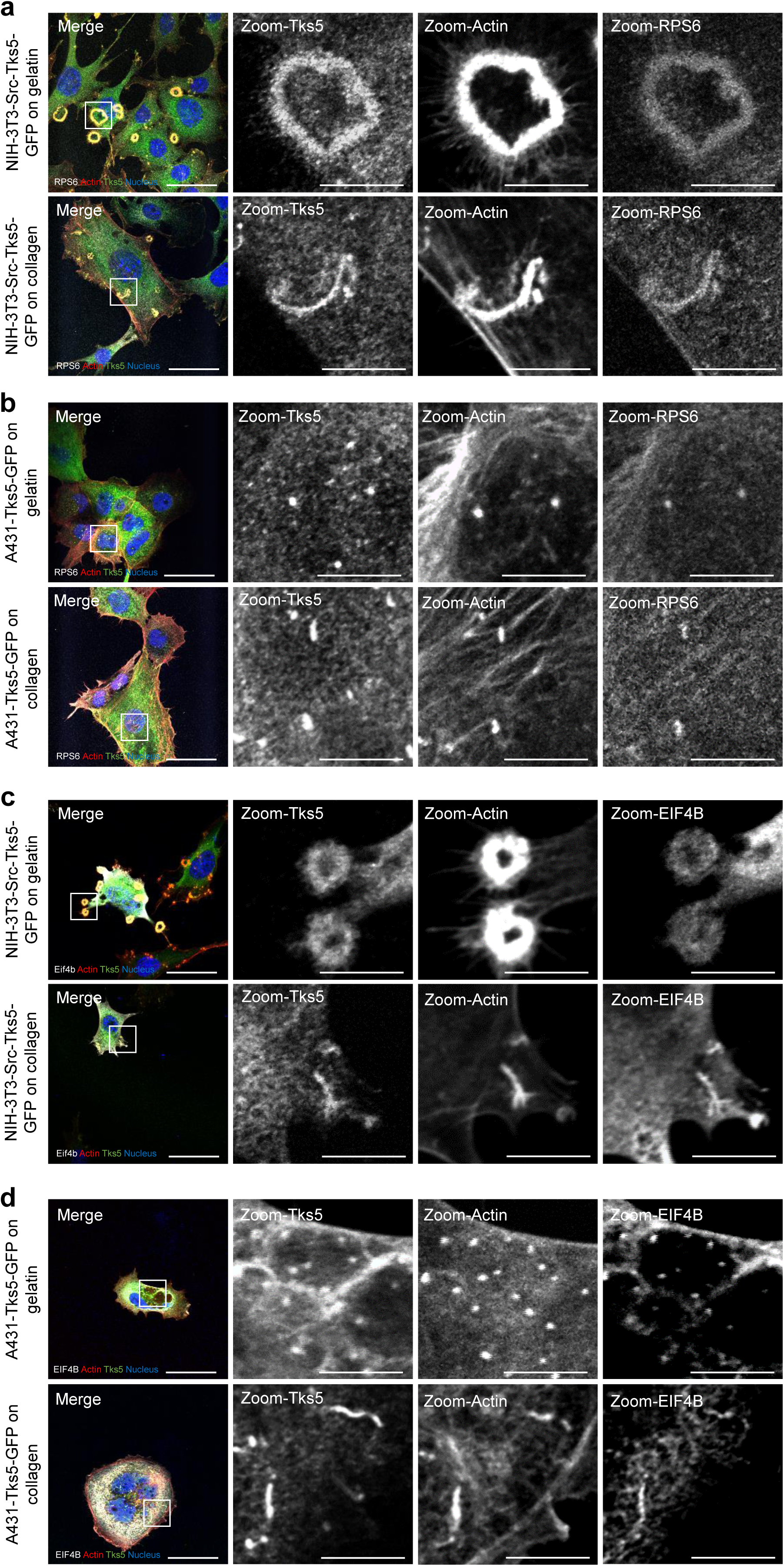
The translation machinery is present in all types of invadosomes. **a)** Confocal microscopy images of NIH3T3-Src-Tks5-GFP cells. The cells were seeded on gelatin or type I collagen and stained for Tks5 in green, actin in red, nuclei in blue and RPS6 in grey. Scale bar: 40µm, zoom: 10µm. **b)** Confocal microscopy images of A431-Tks5-GFP cells. The cells were seeded on gelatin or type I collagen and stained for Tks5 in green, actin in red, nuclei in blue and RPS6 in grey. Scale bar: 40µm, zoom: 10µm. **c)** Confocal microscopy images of NIH3T3-Src-Tks5-GFP cells. The cells were seeded on gelatin or type I collagen and stained for Tks5 in green, actin in red, nuclei in blue and EIF4B in grey. Scale bar: 40µm, zoom: 10µm. **d)** Confocal microscopy image of A431-Tks5-GFP cells. The cells were seeded on gelatin or type I collagen and stained for Tks5 in green, actin in red, nuclei in blue and EIF4B in grey. Scale bar: 40µm, zoom: 10µm.

### Involvement of the translation machinery in invadosome function

In order to investigate and confirm the role of translation in all invadosomes formation and matrix degradation activity from both cell lines, we treated A431 and NIH-3T3-Src cells seeded on gelatin or type 1 collagen with the translation inhibitor cycloheximide (CHX) (Supp Figure 3). Our previous study demonstrated that CHX blocked the formation of rosettes in a short time, after one hour of treatment^21^. By using Western Blot and SUrface SEnsing of translation (SUnSET) assay, we first showed that CHX treatment reduced global translation in all conditions (Supp Figure 3a and 3b). We next confirmed that CHX treatment in NIH-3T3-Src model prevented invadosome rosette formation (Figure 4a, 4b) as previously described^21^. No significant differences for dots and linear invadosomes formation was measured by immunofluorescence quantification between CHX and control group (DMSO) (Supp Figure 3c and 3d). However, CHX treatment limited invadosome degradation activity by A431 and NIH-3T3-Src cells on gelatin and collagen matrices (Figure 4c, 4d). These results showed that global translation is involved in degradation capacity of all invadosomes and formation only in rosette organization.

**Figure 4:**
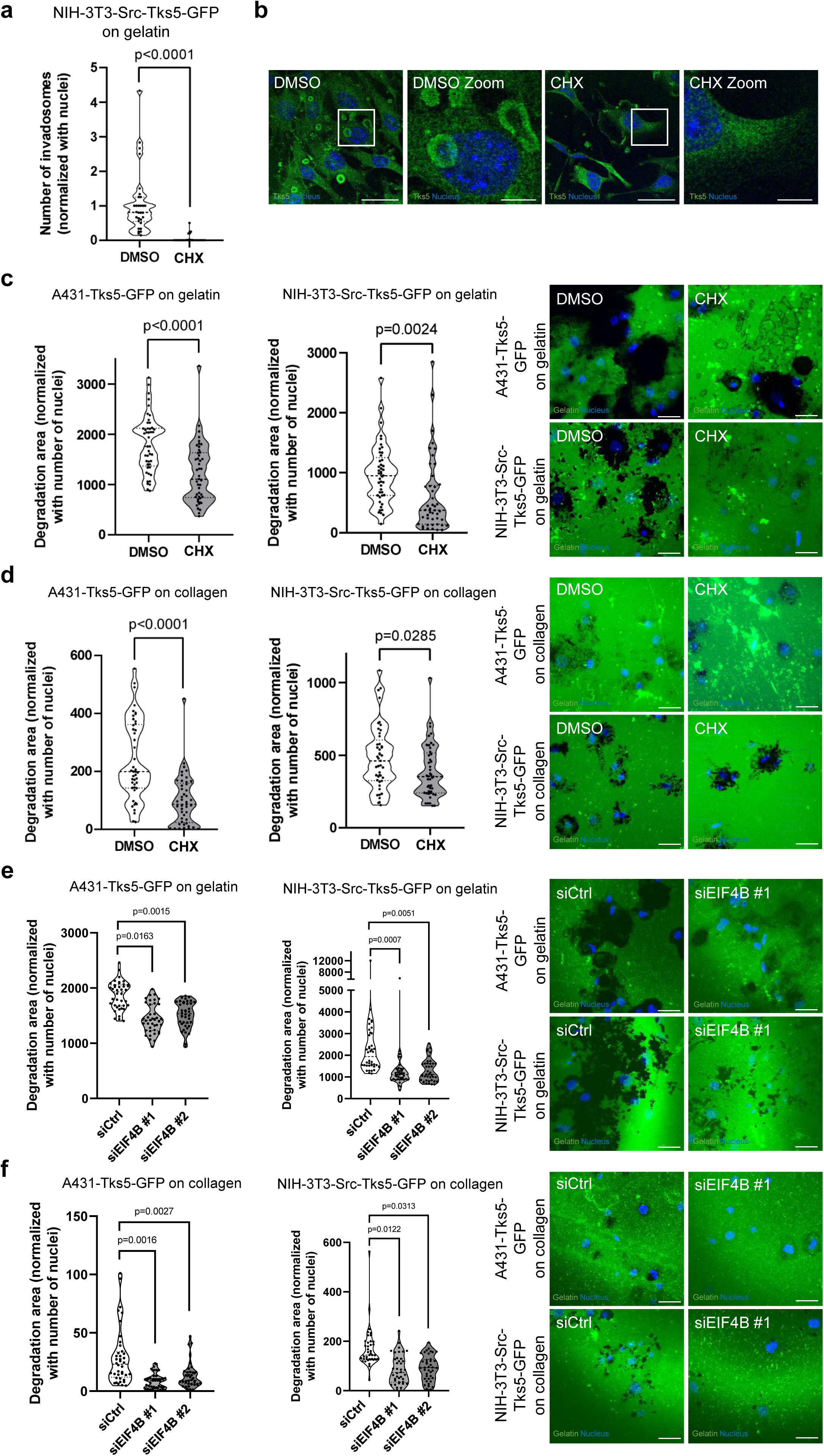
Involvement of the translation machinery in invadosome activity. **a)** Quantification of the numbers of invadosomes per cell treated (CHX) or not (DMSO) with cycloheximide in NIH3T3-Src-Tks5-GFP cells seeded on gelatin. Values represent the mean +/- SEM of n=3 independent experiments (10 images per condition and per replicate) and were analyzed using student t-test. **b)** Representative images of the quantification of invadosome formation in NIH3T3-Src-Tks5-GFP cells seeded on gelatin. Tks5 is stained in green and nuclei in blue. Scale bar: 40µm, zoom: 10µm. **c)** Left: Quantification of the ECM degradation properties of A431 and NIH3T3-Src-Tks5-GFP cells on gelatin treated (CHX) or not (DMSO) with cycloheximide. Values represent the mean +/- SEM of n=3 independent experiments (15 images per condition and per replicate) and were analyzed using student t-test. Right: Representative images of the quantification of ECM degradation of A431 and NIH3T3-Src-Tks5-GFP cells on gelatin. Gelatin is stained in green and nuclei in blue. Scale bar: 40µm, zoom: 10µm. **d)** Left: Quantification of the ECM degradation properties of A431 and NIH3T3-Src-Tks5-GFP cells on collagen treated (CHX) or not (DMSO) with cycloheximide. Values represent the mean +/- SEM of n=3 independent experiments (15 images per condition and per replicate) and were analyzed using student t-test. Right: Representative images of the quantification of ECM degradation of A431 and NIH3T3-Src-Tks5-GFP cells on collagen. Gelatin is stained in green and nuclei in blue. Scale bar: 40µm, zoom: 10µm. **e)** Left: Quantification of the ECM degradation properties of A431 and NIH3T3-Src-Tks5-GFP cells on gelatin treated (siEIF4B#1 and #2) or not (siCtrl) with siEIF4B. Values represent the mean +/- SEM of n=3 independent experiments (15 images per condition and per replicate) and were analyzed using student t-test. Right: Representative images of the quantification of ECM degradation of A431 and NIH3T3-Src-Tks5-GFP cells on gelatin. Gelatin is stained in green and nuclei in blue. Scale bar: 40µm, zoom: 10µm. **f)** Left: Quantification of the ECM degradation properties of A431 and NIH3T3-Src-Tks5-GFP cells on collagen treated (siEIF4B#1 and #2) or not (siCtrl) with siEIF4B. Values represent the mean +/- SEM of n=3 independent experiments (15 images per condition and per replicate) and were analyzed using student t-test. Right: Representative images of the quantification of ECM degradation of A431 and NIH3T3-Src-Tks5-GFP cells on collagen. Gelatin is stained in green and nuclei in blue. Scale bar: 40µm, zoom: 10µm.

As we identified EIF4B in all invadosomes (Figure 2c; Supp Figure 2c), we assessed the role of EIF4B in invadosome degradation activity. By using small interfering RNA (siRNA) strategy we silenced EIF4B expression in both cell lines (Supp Figure 3e) and demonstrated that EIF4B depletion prevented invadosome rosette formation in NIH-3T3-Src model and linear invadosome formation in A431 model (Supp Figure 3f). However, EIF4B depletion reduced matrix degradation activity in all organizations of invadosomes from both cell lines (Figure 4e, 4f). These results showed that EIF4B is involved in degradation capacity of all invadosomes.

These data confirm the involvement of the translation and more specifically of the EIF4B protein in the matrix degradation activity of all invadosomes.

### Recruitment of ER into invadosome rosettes

In addition to translation proteins, the mass spectrometry analysis highlighted the presence of ER-related proteins such as RTN4, LRRC59 or RRBP1 in all invadosomes linked with Tks5 (Figure 2c). Ribosomes and ER proteins have previously been identified in the proteome of invadosome rosettes^21^.

In order to assess the involvement of the ER in invadosome formation and function we used the rosette model, which is the largest invadosome structure, facilitating analysis and imaging. We used lifeact-mRuby-expressing NIH-3T3-Src cells transfected with KDEL-GFP to detect ER presence. By video microscopy, we showed that the ER was recruited at the level of the rosette in formation and that there was an enrichment of the ER inside the actin core, as previously described in fixed cells^21^ (Figure 5a, Video 1). Furthermore, time-lapse imaging clearly shows that the RE is not present in all the cytoplasm and revealed that the ER forms “arms” leading to a “ER flow” heading towards the forming rosette then accumulating within the actin core (Figure 5a, Vidéo 1). The transversal sections confirm the enrichment as we can clearly see that the ER arm is inserted within the rosette actin core (Figure 5a). 3D reconstruction allowed us to visualize the accumulation of the ER within the rosette core (Figure 5b). In order to confirm the presence of ER at the level of rosettes, we performed correlative light and electron microscopy (CLEM). Correlation between actin fluorescent images and electron microscopy allowed the identification of ER tubules associated with the rosettes (Figure 5c).

**Figure 5:**
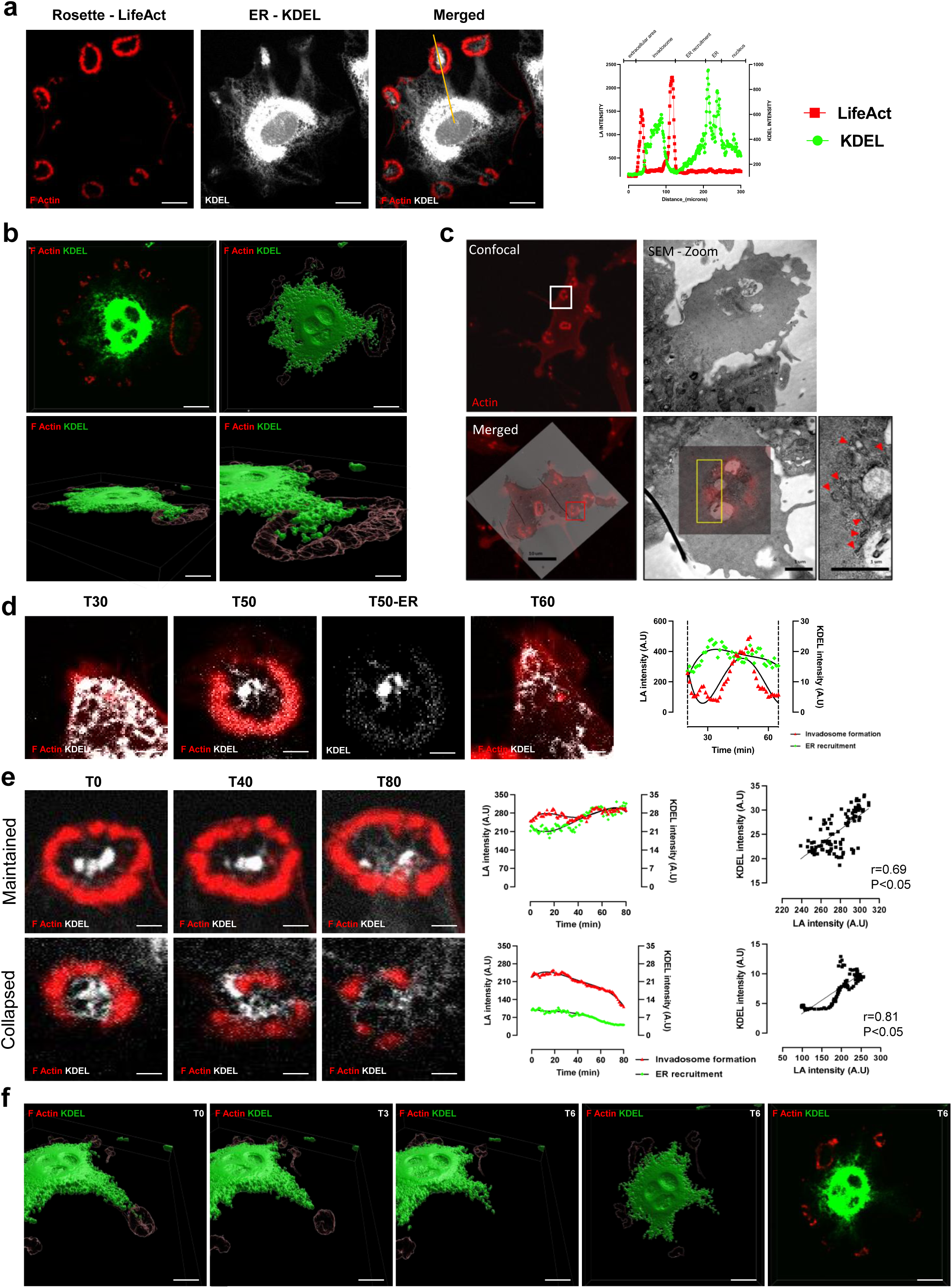
Recruitment of ER into invadosome rosettes. **a)** Representative images from time-lapse video microscopy of lifeact-mRuby (red)-expressing NIH-3T3-Src cells transfected with KDEL-GFP (white). The lifeact-mRuby and KDEL-GFP were measured along the yellow axis. Scale bar: 20µm. **b)** Projection of a z-stacks and Imaris surface 3D renderings are shown, visualizing that KDEL-GFP (green) localizes inside invadosome during maturation. Scale bar: 20µm. **c)** CLEM images of invadosomes of NIH-3T3-Src cells on gelatin matrix. Actin is stained in red. Red arrows show the endoplasmic reticulum. Scale bar: 10µm and 1µm. **d)** Representative images from time-lapse video microscopy of lifeact-mRuby (red)-expressing NIH-3T3-Src cells transfected with KDEL-GFP (white). Intensity level of lifeact-mRuby and KDEL-GFP were measured in initiation step in rosette. Scale bar: 5µm. **e)** Representative images from time-lapse video microscopy of lifeact-mRuby (red)-expressing NIH-3T3-Src cells transfected with KDEL-GFP (white). Intensity level of lifeact-mRuby and KDEL-GFP were measured in maintained and collapsed in rosettes. Scale bar: 5µm. **f)** Projection of a z-stacks and Imaris surface 3D renderings are shown, visualizing that KDEL-GFP (green) escape from invadosome during his collapse. Scale bar: 20µm.

In order to determine the dynamic of ER recruitment during invadosome formation, time-lapse imaging was performed. The analysis revealed that ER recruitment occurs before actin polymerization and formation of rosette-like invadosomes (Figure 5d, Video 2). In addition, we showed that the maintenance of the structures requires the presence of ER (Figure 5e upper panel, Video 3a) while a drop in ER recruitment is associated with a collapse of the structure (Figure 5e lower panel, Video 3b). 3D reconstruction confirmed that the collapse of the rosette structure is associated with a withdrawal of the ER within the structure with the “ER flow” heading towards the interior of the cell (Figure 5f). These results therefore confirm the presence but also a potential involvement of the ER in the rosette formation and maintenance over time.

### Involvement of ER-associated machinery translation in invadosome formation

As we identified the presence and the determinant role of translation and ER in invadosome formation, we investigated the role of ER-associated translation in invadosome formation and matrix degradation activity. Previously published results showed the involvement of the ER in the formation and function of invadosomes via the involvement of the protrudin protein or by the involvement of the calnexin/erp57 protein complex respectively^32,33^. To determine the role of ER-associated translation in the dynamic of invadosome rosettes formation, we performed a time-lapse experiment with cells treated with CHX to inhibit translation (Figure 6a, Video 4). Translation inhibition led to the collapse of the rosette structure while the ER was still present at the ‘previous invadosome site” without re-formation of invadosomes (even if some actin polymerization events were recorded) (Figure 6a, Video 4). This result suggests that ER-associated translation is mandatory for invadosome persistence.

**Figure 6:**
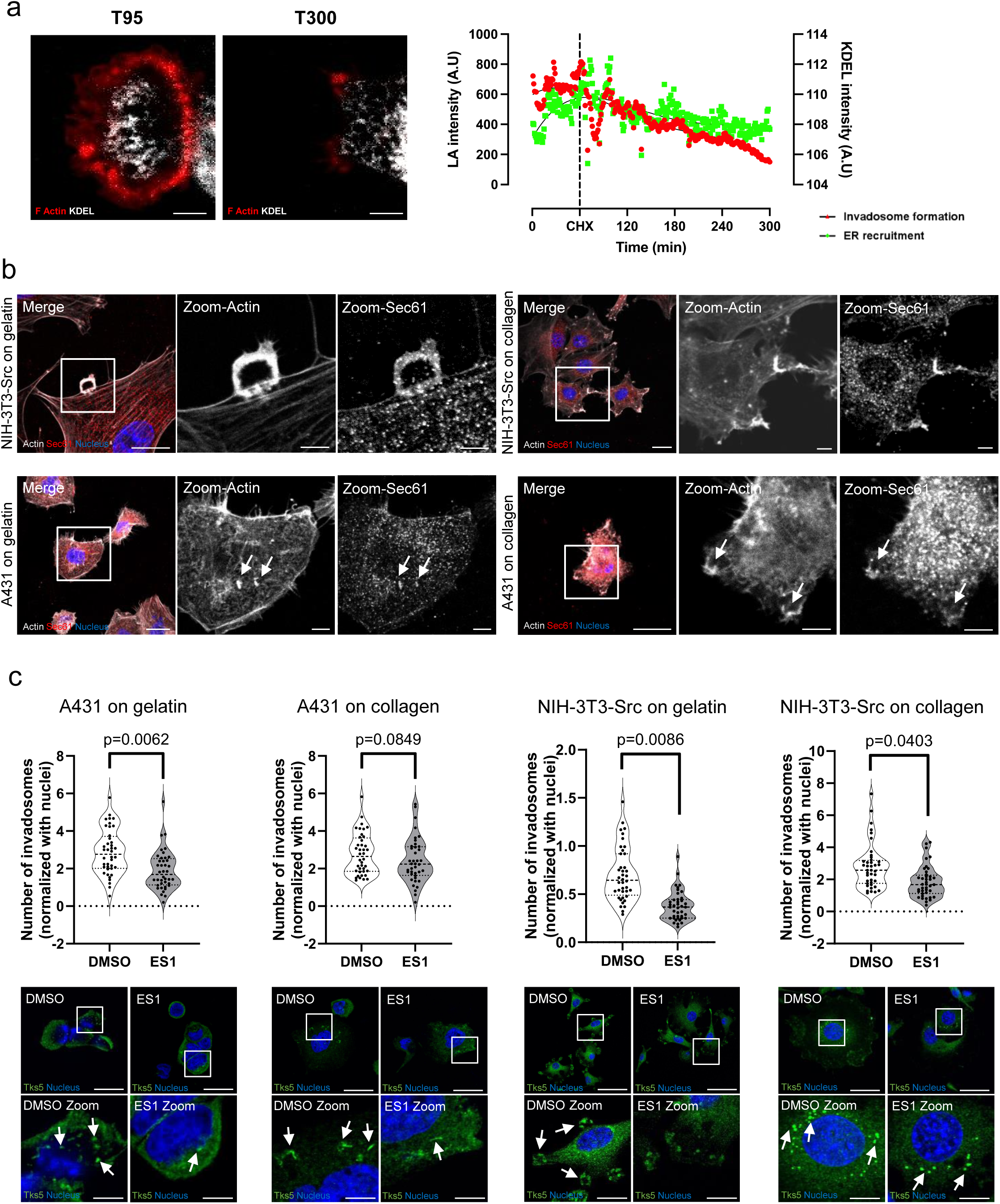
Involvement of ER-associated machinery translation in invadosome formation. **a)** Lifeact-mRuby and KDEL-GFP signals were recorded in NIH-3T3-Src cells treated with cycloheximide (CHX; 35µM) at T60 min. Scale bar: 5 µm. **b)** Confocal microscopy images of NIH-3T3-Src and A431 cells. The cells were seeded on gelatin or type I collagen and stained for Sec61 in red, nuclei in blue and Actin in grey. Scale bar: 20µm, zoom: 5µm. **c)** Upper part: Quantification of invadosome formation in A431 and NIH-3T3-Src cells seeded on gelatin or type 1 collagen treated or not (DMSO) with Sec61 translocon inhibitor (ES1) during one hour. Lower part: Representative images of the quantification of invadosomes formation in A431 and NIH3T3-Src-Tks5-GFP cells seeded on gelatin or type 1 collagen treated or not (DMSO) with ES1. The arrows show the invadosomes. Tks5 is stained in green and nuclei in blue. Scale bar: 40µm, zoom: 10µm.

To further characterize the involvement ER-associated translation in invadosome formation, we treated cells with Sec 61 translocon inhibitor (ES1) which blocks the translocation of proteins to the ER. Indeed, Sec61 is a well-described ER marker that allows protein insertion into the ER but also is a key player for ribosomes docking to the ER^34^. Furthermore, Sec61 was identified by our mass spectrometry analysis for A431 cells on plastic and collagen (Supp Table 1) and localized with invadosome structures (Figure 6b). ES1 treatment on A431 and NIH-3T3-Src cells seeded on gelatin or type 1 collagen matrix led to a decreased number of all organizations of invadosomes (Figure 6c). These results therefore suggest that the translation machinery must be adapted somewhat depending on the matrix context. This would explain why the translation machinery is only involved in the degradation function for dots and linear invadosomes while it is involved in the formation and degradation function for rosettes, which are larger structures.

All together, these results demonstrate that the ER-associated translation machinery identified in Tks5 interactome is an essential element to maintain the formation of all types of invadosomes.

## Discussion

We provided here a molecular characterization of all types of invadosomes by identifying the interactome of the common and universal invadosome marker Tks5. We used co-immunoprecipitation coupled with mass spectrometry to identify new proteins shared by different invadosome types such as dots, rosettes and linear invadosomes (Figure 1). This strategy allowed us to identify proteins already known in the literature but also to identify new ones. We showed that Tks5 is linked to translation proteins and confirm that translation is involved in the degradation function in all types of invadosomes, including linear invadosomes.

Indeed, we identified 46 translation proteins with some already known in the literature to be involved in invadosome activity such as IGF2BP2, IGF2BP3 and FGD1^35–38^. Regarding the other proteins identified and already described in invadosomes, microtubule-associated proteins and especially MAP4 and LAMP1 were described in podosomes^24,27,39^ and ITGA5 was described in invadopodia^31^. PTBP1 was also identified in podosomes as participating in their formation^40^. We further demonstrated here that the matrix receptor CD44 which was previously described in podosomes and invadopodia but not in linear invadosomes^28,41^ is also a common marker of invadosomes (Figure 2). These known proteins allowed us to serve as quality control for mass spectrometry experiments.

Thanks to mass spectrometry experiments, we were also able to validate the presence of translation proteins in linear invadosomes. We identified the presence of EIF4B and RPS6 proteins in all invadosome organizations (Figure 3). Interestingly, these two proteins are known in the literature to be targeted by the mTOR/ S6K1 pathway. Indeed, S6K1 is be able to directly phosphorylate EIF4B to allow the recruitment of EIF4A and EIF3 as well as the RPS6 protein to initiate translation^42^. While we demonstrated that global translation is mandatory for degradation activity induced by invadosomes, the inhibition of global translation did not prevent invadosome formation (excepted for rosettes) (Figure 4). Nevertheless, we also showed that ER-associated translation regulates all type of invadosomes formation suggesting that ER has a major role in invadosome formation and the associated degradation activity. We hypothesize that ribosome docking to ER, through Sec61, enhances protein synthesis of determinant elements for invadosome formation. A recent study demonstrated that an early increase of Sec61 enhanced the expression of pro-invasive proteins, followed by an expansion of the ER and the Golgi, leading to the delivery of these pro-invasive proteins to the cell membrane^43^.

In addition to the translation proteins identified, proteins present in the ER have also been highlighted by our proteomic analysis such as RTN4 or LRRC59. Since ER is also involved in protein translation, this suggested the presence of a complex translation mechanism within all invadosomes. We showed that the ER is recruited at the rosette, so enrichment of the ER could be linked to the rosette functioning properly, notably for its degradation capacity. In addition, our data show that removal of the ER leads to a collapse of the rosette structure suggesting a link between ER and invadosome formation and degradation activity. Furthermore, we showed that the ER was necessary for initiating and maintaining the invadosome rosettes (Figure 5 and 6). Indeed, the spatiotemporal aspect of invadosome formation in the initiation, stabilization or collapse phases is a major question^44,45^. Here, we show that the ER plays a main role in these different stages.

In literature, Pedersen et *al.* also showed the involvement of ER in invadosome formation: protrudin mediated ER-endosome contact site promotes cell invasion through the translocation of MT1-MMP endosome to the plasma membrane leading to invadopodia outgrowth and degradation^32^. Moreover, one of our recent study showed that two ER proteins, calnexin and ERp57, can be trafficked to invadosomes and are essential for ECM degradation^33^. These results therefore suggest the presence of a complex allowing invadosomes to have their own translation machinery to facilitate cell invasion with local translation of specific mRNA^21,46^. Tks5 could therefore act as a molecular crossroads allowing translational proteins to cluster in the same place, leading to rapid localization and greater efficiency for the cell in the formation and function of invadosomes.

In order to better understand the link between the translation and the ER in all types of invadosomes, we first targeted the EIF4B protein, identified by mass spectrometry. Using siRNA strategy, we showed that EIF4B is required in degradation function. Recent studies already showed the involvement of EIF4B in invasion. Indeed, Cao and al. showed that the pathway PIM1/EIF4B/c-MET pathway was involved in tumor migration and invasion in lung adenocarcinoma^47^. In the same way, the GMAN/p-EIF4B pathway leads to cell proliferation and invasion in hepatocellular carcinoma (HCC)^48^.

We also identified many proteins associated with EIF3 and EIF4 complexes such as EIF3C, EIF4A3, EIF4E or EIF3H in at least three out of four types of invadosomes. Similarly, these complexes are already known in the literature to participate in the process of cell invasion.

The YTHDF1/EIF3C axis is involved in the invasion process in ovarian cancer^49^, EIF3H participates to the invasion of hepatocellular carcinoma^50^ and the EIF4A3/FLOT1 pathway promotes invasion and tumor proliferation in lung adenocarcinoma^51^. These results suggest the involvement of these complexes in the degradation mechanism that could be common to all types of invadosomes.

Altogether, these results provide new knowledge specifically into invasive structures and on the invasion process more broadly. Indeed, we reported new molecular components common to all types of invadosomes through the identification of Tks5 interactome. This thus provides a better understanding of the mode of operation of invadosomes, in particular via the involvement of the ER-associated translation machinery which is involved into invadosome formation.

### Material and methods

#### Antibodies and reagent

Anti-GFP (catalog number ab6673) antibody was purchased from Abcam. Anti-Tks5 (G-7, catalog number sc-376211), and anti-CD44 (DF1485, catalog number sc-7297) were purchased from Santa Cruz Biotechnology, Inc. Anti-MAP4 (AG1741, catalog number 11229-1-AP) was purchased from Proteintech. Anti-RPS6 (S369, catalog number AP22299a) was purchased from Abgent. Anti-DDK (OTI2F7, catalog number TA500288) was purchased from OriGene Technologies, Inc. Anti-EIF4B (catalog number 3592) was purchased from Cell Signaling Technology. Secondary antibodies for immunofluorescence IRDye® 680RD Conjugated Goat anti-mouse (926-68070, LI-COR Biotech.), IRDye® 680RD Conjugated Goat anti-rabbit, (926-68071, LI-COR Biotech.), IRDye® 800CW Conjugated Goat anti-mouse (926-32210, LI-COR Biotech.), IRDye® 800CW Conjugated Goat anti-rabbit (926-32211, LI-COR Biotech.). The Atto 647N conjugated Goat anti-mouse-IgG was purchased from Sigma-Aldrich (catalog number 50185). DAPI (catalog number H21486), phalloidin (catalog number FP-BZ-9630) were purchased from Interchim. Type I collagen from rat tail(catalog number 354236) was purchased from Corning. Sec61 translocon was inhibited by pharmacological approach using Eeyarestatin 1 (ES1, catalog number E1286 Sigma-Aldrich) at 5µM during 1 hour. Species-specific fluorescent far-red coupled secondary antibody for western blot IRDye 680CW goat (polyclonal) anti-rabbit IgG (H+L) (catalog number 926-68070) or IRDye 800CW goat (polyclonal) anti-mouse IgG (H+L) (catalog number 926-32210) were purchased from LI-COR Biotech. GM6001 MMP Inhibitor (catalog number CC10), Cycloheximide (CHX) (catalog number 01810) and Manganese(II) (catalog number M7899) were purchased from Sigma-Aldrich.

#### Cell culture

The A431 cell line was generously donated by Dr. Julie Dechanet-Merville (CNRS 5164, Bordeaux) and authenticated. NIH-3T3-Src cells were generously donated by Dr. Sara Courtneidge (Université of Portland, USA). A431 and NIH-3T3-Src cells were maintained in DMEM high glucose (Gibco) supplemented with 10% fetal bovine serum (FBS, 139 Sigma Aldrich) at 37°C in a 5% CO2 incubator. In this study, we used A431 and NIH-3T3-Src cells transduced with a lentiviral plasmid expressing Tks5 fused to GFP, and the cell line was named A431-Tks5-GFP and NIH3T3-Src-Tks5-GFP.

### Lentiviral transduction

A431 and NIH-3T3-Src cells were seeded in a 6-well plate, then transduced after 24 h at a multiplicity of infection of 10 μg, either with a lentiviral plasmid containing Tks5 couple to GFP, or with the GFP alone plasmid encoding as a control . The transduced cells were selected using puromycin treatment at 1µg/mL. Co-transduction of lifeact-mRuby and KDEL-GFP NIH-3T3-Src cells has been performed as described above but double positive cells were selected and sorted using BD FACSAria Cell Sorter (BD Bioscience).

#### Transfection

EIF4B-DDK plasmid was purchased from OriGene Technologies, Inc. This plasmid was transfected (2,4 µg) using JetPrime (PolyPlus Transfection) following the manufacturer’s instructions. siRNA oligonucleotides (20 nM) targeting eIF4B (Human: s4573 & s4574; Mouse: s93804 & s93805 ThermoFisher) were transfected using Lipofectamine RNAiMax (Invitrogen) according to the manufacturer’s instructions.

#### Coating of coverslips with extracellular matrix (ECM)

Cells were seeded on gelatin coverslips with or without coating of collagen. Type I collagen (rat tail, catalog number 354236, Corning) was diluted in DPBS containing calcium and magnesium (Dulbecco’s Buffered-Phosphate Saline, Gibco) at a final concentration of 0.5mg/ml and polymerized for 4 h at 37°C. Gelatin-coated coverslips were made using gelatin from Sigma-Aldrich (catalog number G1393). Coverslips were incubated with gelatin for 20 min before and then fixed with 0.5% glutaraldehyde (Electron Microscopy Science) for 40 min at room temperature. Coverslips were washed 3 times with PBS before cell seeding.

#### Immunofluorescence and imaging

Cells were fixed using 4% paraformaldehyde (PFA) (Electron Microscopy Sciences) for 10 min at room temperature and then rinsed 3 times with PBS. Cells were permeabilized using 0.2% Triton X-100 (catalog number T9284, Sigma-Aldrich) for 10 min at room temperature before being rinsed twice with PBS. Cells were then incubated with primary antibodies (1:100 diluted in PBS-4%BSA) for 40 min at room temperature, rinsed 3 times with PBS. The cells were then incubated with secondary antibodies (1:200 diluted in PBS-4%BSA) for 30 min at room temperature. Nuclei were stained with DAPI (1:1,000 dilution) and actin was stained with phalloidin (1:200 dilution). Coverslips were mounted on microscope slides using Fluoromount-G mounting media (catalog number 0100-01, SouthernBiotech) and were imaged using SP5 confocal microscope (Leica, Leica Microsystems GmbH, Wetzlar, Germany). Collagen is observed by internal reflection microscopy (IRM). Images were analyzed using ImageJ or Fiji softwares.

#### Live-cell imaging

Cells were seeded on glass bottom dishes then image acquisitions were performed every 2 min for 4 using LiveSR spinning-disk microscope (Leica, Leica Microsystems GmbH, Wetzlar, Germany) with environmental control at 37°C and CO_2_ provider. Z-stacks were performed then deconvolution imaging were used for 3D reconstruction (Imaris software).

#### Correlative light and electron microscopy (CLEM)

For performing CLEM of invadosome rosettes, we did micro-pattern on ACLAR^®^ films (Catalog number 77850-12, EMS) by the laser microdissection^52^. Solution of gelatin (0.5 mg/mL) was pre-coated on these patterned ACLAR^®^ substrates in a 24-well plate. The lifeact-mRuby expressed NIH-3T3-Src cells were then seeded at 20,000 cells/mL on ACLAR^®^ films and incubated overnight. The film was transferred to a 35mm glass bottom dish (Catalog number P35G-1.0-20-C, MatTek) for confocal imaging (Leica SP2) using a × 63 objective (NA:1.32, Leica Microsystems). Position of the region of interest (ROI) according to the landmarks visible in transmitted light. High-magnification images of the cells and rosettes of interest were acquired as well as low-magnification images. After imaged, cells were fixed in primary fixation buffer contained 2.5% PFA (Catalog number 15713, EMS) and 2.5% GA (Catalog number 16220, EMS) in 0.1M cacodylate buffer (Catalog 11652, EMS) for 1 h at room temperature. Post-fixation was performed in 1% osmium tetroxide (Catalog number 19150, EMS) in 0.1M cacodylate buffer for 1 h on ice. Cells were stained with 2% uranyl acetate, and then dehydrated in sequential gradient alcohol baths and infiltrated with Epon resin. We then inversed the film on the top of the plastic embedding capsule (Catalog number 69910-05, EMS) following overnight polymerization at 60 °C thus embedding the intact cell monolayer of adhesion cells. The following day, the capsules were filled with fresh Epon and polymerized again overnight at 60 °C. Accurate positioning of the ROI can be relocated by a stereomicroscope according to the micro-pattern. Finally, the precise trimmed resin block was done serial sectioning (thickness: 70 nm) and all the sections were collected on slot grids for TEM observation. Micrographs were obtained at transmitted electron microscope (CM12, FEI company) with a CCD camera (ORIUS, Gatan). For the image process, overlay the fluorescent and the EM images in Fiji, and manually rescale or rotate the data until the cell boundaries align.

#### Western Blot

Cells were washed once with cold PBS on ice, and then were scraped with RIPA lysis buffer containing phosphatase and protease inhibitor cocltails (catalog number 04906845001, Merck) and lysed for 30 min on ice. Cell lysates were clarified by centrifugation at 13,000g for 15 min at 4°C. Protein concentrations were determined using BCA reagent (DC^TM^ Protein Assay, BioRad), according to the manufacturer’s protocol. Lysates were heat at 95°C for 10 min and then separated by SDS-PAGE electrophoresis on 10% acrylamide gels (FastCast™ Acrylamide kit, BioRad) at 110V for 60 min. Samples were transferred onto nitrocellulose membranes using a Transblot transfer system (BioRad) and membranes were blocked using 5%BSA with0.2%Tween (catalog number P7949, Sigma-Aldrich) in TBS buffer (catalog number ET220, Euromedex) for 1 h at room temperature before being incubated with primary antibodies (1:1,000 dilutiond in blocking buffer as described above) overnight at 4°C. The next day, membranes were washed 3 times with TBST and incubated with secondary antibodies (1:5,000 dilution with 5% BSA inTBST) for 30min at room temperature. Membranes were washed 3 times with TBST before exposure using a Chemidoc system (BioRad).

#### Sunset assay

Cells were treated with puromycin (10mg/ml) during 10min at 37°C then washed twice in ice-cold PBS for protein extraction as described above in Western Blot section. For negative control we pre-treated cells with the translation inhibitor cycloheximide (35mM) during 10min at 37°C.

#### Immunoprecipitation

For immunoprecipitation experiments, 4.5 million cells were necessary and technical triplicate were performed. NIH-3T3-Src-Tks5-GFP and A431-Tks5-GFP cells were seeded on plastic or in dishes with a fibrillar collagen I matrix. Lysates were centrifuged for 15 min at 13,000g at 4°C and supernatants were immunoprecipitated with the GFP-Trap®_MA kit (Goldstandard) according to the supplier(s) recommendations. The immunoprecipitates were washed three times in lysis buffer, and then assayed using the Lowry method (DC^TM^ Protein Assay kit, Biorad) using a spectrophotometer at 750nm.

#### Mass spectrometry

The steps of sample preparation and protein digestion by the trypsin were performed as previously described^53^. NanoLC-MS/MS analysis were performed using an Ultimate 3000 RSLC Nano-UPHLC system (Thermo Scientific, USA) coupled to a nanospray Q Exactive Hybrid Quadruplole-Orbitrap mass spectrometer (Thermo Scientific, USA). Each peptide extracts were loaded on a 300 µm ID x 5 mm PepMap C_18_ precolumn (Thermo Scientific, USA) at a flow rate of 20 µL/min. After a 5 min desalting step, peptides were separated on a 75 µm ID x 25 cm C_18_ Acclaim PepMap® RSLC column (Thermo Scientific, USA) with a 4-40% linear gradient of solvent B (0.1% formic acid in 80% ACN) in 108 min. The separation flow rate was set at 300 nL/min. The mass spectrometer operated in positive ion mode at a 1.8 kV needle voltage. Data were acquired using Xcalibur 3.1 software in a data-dependent mode. MS scans (m/z 350-1600) were recorded at a resolution of R = 70000 (@ m/z 200) and an AGC target of 3 x 10^6^ ions collected within 100 ms. Dynamic exclusion of selected precursors was set to 30 s and top 12 ions were selected from fragmentation in HCD mode. MS/MS scans with a target value of 1 x 10^5^ ions were collected with a maximum fill time of 100 ms and a resolution of R = 17500. Additionally, only +2 and +3 charged ions were selected for fragmentation. Other settings were as follows: no sheath and no auxiliary gas flow, heated capillary temperature, 200°C; normalized HCD collision energy of 27% and an isolation width of 2 m/z. Protein identification and Label-Free Quantification (LFQ) were done in Proteome Discoverer 3.0. CHIMERYS node using prediction model inferys_2.1 fragmentation was used for protein identification in batch mode by searching against a Uniprot *Homo sapiens* database (82 439 entries, release March 2023) or a UniProt Mus musculus database (55 029 entries, release March 2023). Two missed enzyme cleavages were allowed for the trypsin. Peptide lengths of 7-30 amino acids, a maximum of 3 modifications, charges between 2-4 and 20 ppm for fragment mass tolerance were set. Oxidation (M) and carbamidomethyl (C) were respectively searched as dynamic and static modifications by the CHIMERYS software. Peptide validation was performed using Percolator algorithm^54^ and only “high confidence” peptides were retained corresponding to a 1% false discovery rate at peptide level. Minora feature detector node (LFQ) was used along with the feature mapper and precursor ions quantifier. The normalization parameters were selected as follows: (1) Unique peptides, (2) Precursor abundance based on intensity, (3) No normalization was applied, (4) Protein abundance calculation: summed abundances, (5) Protein ratio calculation: pairwise ratio based and (6) Missing values are replaced with random values sampled from the lower 5% of detected values. Quantitative data were considered for master proteins, quantified by a minimum of 2 unique peptides and a fold changes ≥ 2.

#### Inhibition of translation machinery present in invadosomes

A431-Tks5-GFP and NIH-3T3-Src-Tks5-GFP cells were seeded on coverslips coated with gelatin (dots/rosettes) or gelatin and type I collagen (linear invadosomes). The cells were treated with 35µM of translation inhibitor cycloheximide (CHX) (01810, Sigma-Aldrich) or DMSO (control) (D8418, Sigma-Aldrich) 2 h after seeding. For cells seeded on collagen, cells were fixed 4h after seeding. For cells seeded on gelatin, cells were fixed 24 h after seeding by using 4% PFA. The immunofluorescence was performed as previously described. A total of 10 images were performed for each condition using a confocal microscope (SP5, Leica) and experiments were done in three biological replicates. The number of invadosomes were quantified using ImageJ and normalized to the number of nuclei. For Western Blot analysis, cells were seeded on plate and treated with 35µM of cycloheximide (CHX) or DMSO (control) 2h after seeding. Before extraction, cells were treated with Puromycin at a concentration of 10µg/mL. Western Blot was performed as previously described.

#### Image analysis of fluorescence intensities and area of gelatin degradation

Sterile coverslips were coated with Oregon Green™ 488 conjugated gelatin 488 (catalog number G13186, Thermofisher) for 20 min and fixed with 0.5% glutaraldehyde (catalog number 15960, EMS) for 40 min. Then, the same experiment as described before was performed. Coverslips were imaged under the epifluorescence microscope (Zeiss) using the x 63 oil immersion objective. A total of 15 images were acquired for each condition, and experiments were done in three biological replicates. The area of degradation was quantified using ImageJ and normalized to the number of nuclei in each image.

#### Statistical analysis

Data are reported as the mean ± SEM of at least three experiments. Statistical significance (*p* < 0.05 or less) was determined using analysis of variance (ANOVA) (one way, two way followed by a Bonferroni’s correction), unpaired or paired student t-test as appropriate and performed with GraphPad Prism software. Significance levels are shown as p-values.

#### Data and material availability

The mass spectrometry proteomics data have been deposited to the ProteomeXchange Consortium via the PRIDE^55^ partner repository with the dataset identifiers PXD046512 (NIH cells) and PXD046515 (A431 cells).

## Abbreviations

CHX: cycloheximide
ECM: extracellular matrix
EIF4B: eukaryotic translation initiation factor
4B ER: endoplasmic reticulum
RPS6: ribosomal subunit S6

## Acknowledgments

The microscopy was done in the Bordeaux Imaging Center a service unit of the CNRS-INSERM and Bordeaux University, member of the national infrastructure France BioImaging supported by the French National Research Agency (ANR-10-INBS-04). The help of Magali Mondin is acknowledged. We are also grateful to platform of the unit TBMCore (US 005), Oncoprot and FACSility (US 005). We also thank Kuang-Jing Huang from U1109 for his help.

## Author contributions

L.N and B.B designed and performed experiments. S.D.T, J.W.D and A.A.R performed and interpreted mass spectrometry analysis. L.M and J.G contributed to CLEM analysis. L.N, B.B, V.M, E.H and FS wrote the manuscript, F.S designed and led the study.

## Data Availability

All data supporting the findings of this study are available from the corresponding author upon reasonable request. The mass spectrometry data are available via ProteomeXchange with the identifier PXD046512 (NIH cells) and PXD046515 (A431 cells). Source data are provided with this paper.

## Funding

Léa Normand was supported by a PhD scholarship from the SIRIC BRIO and the Région. Benjamin Bonnard was supported by the Inca. This work was supported by Fondation Ruban rose, Inca (PLBIO-2020-122) and Ligue contre le cancer (équipe labellisée 2023-DN/IP/IQ C:17691).

---

**Supp Figure 1:**
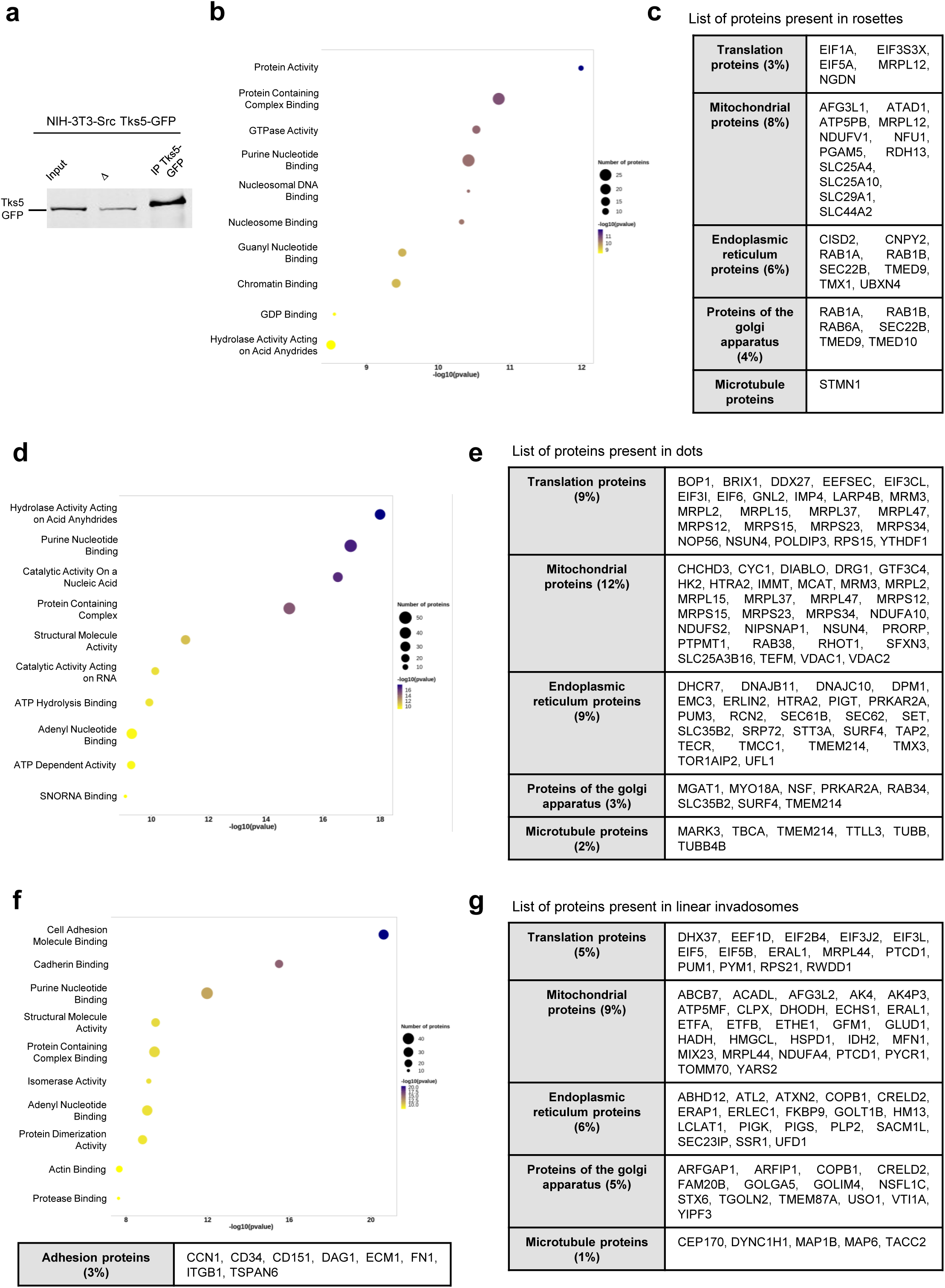
Specific molecular proteins of each invadosomes organization. **a)** Western Blot analysis of the co-immunoprecipitation performed with anti-GFP antibodies in NIH3T3-Src-Tks5-GFP cells. The input, the Δ and the co-IP (IP-Tks5-GFP) fractions are shown. **b)** Bubble plot of the proteins related pathway in rosettes. **c)** Classification table of the specific proteins identified in rosette. Proteins are grouped into five main categories corresponding to translation proteins, mitochondrial proteins, ER proteins, proteins of the Golgi apparatus and microtubules proteins. **d)** Bubble plot of the proteins related pathway in dots. **e)** Classification table of the specific proteins identified in dots. Proteins are grouped into five main categories corresponding to translation proteins, mitochondrial proteins, ER proteins, proteins of the Golgi apparatus and microtubules proteins. **f)** Bubble plot of the proteins related pathway in linear invadosomes and classification table of the adhesion proteins specific to linear invadosomes organizations. **g)** Classification table of the specific proteins identified in linear invadosomes. Proteins are grouped into five main categories corresponding to translation proteins, mitochondrial proteins, ER proteins, proteins of the Golgi apparatus and microtubules proteins.

**Supp Figure 2:**
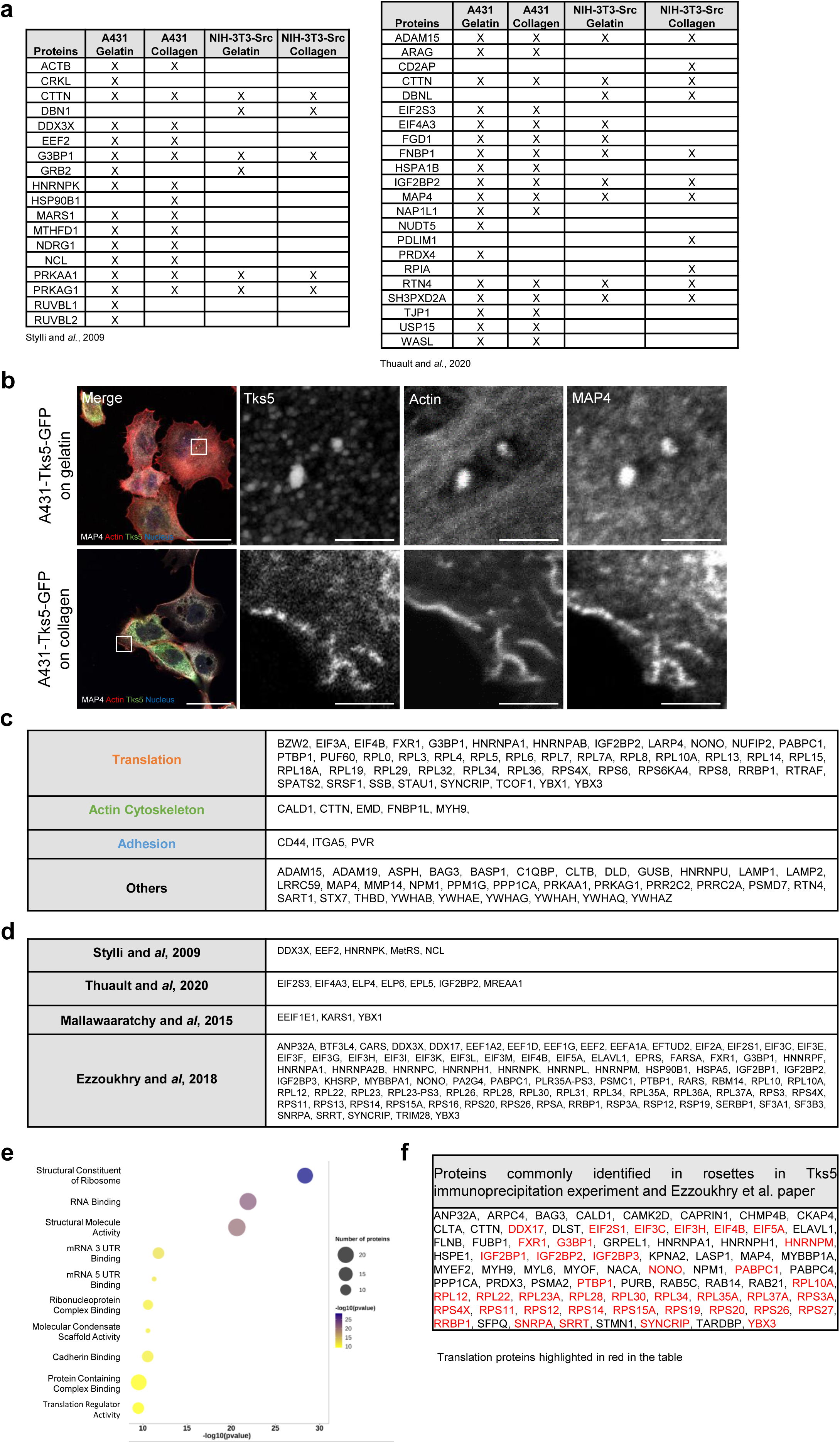
Characterization and validation of proteins identified by mass spectrometry as partner of Tks5. **a)** Table of proteins identified by mass spectrometry in at least one type of invadosomes and already identified in the literature as partner of Tks5 in 293T cells (Stylli and *al.*) and MDA-MB-231 cells (Thuault and *al.*). **b)** Confocal microscopy images of A431-Tks5-GFP cells. The cells were seeded on gelatin or type I collagen and stained for Tks5 in green, actin in red, nuclei in blue and MAP4 in grey. Scale bar: 40µm, zoom: 5µm. **c)** Classification table of the 88 common proteins identified. Proteins are grouped into three main categories corresponding to translation, adhesion and actin cytoskeleton. **d)** Table of translation proteins already identified in invadosomes in the literature **e)** Bubble plot of proteins related pathway identified by mass spectrometry thanks to Tks5 and present in data already published by Ezzoukhry and *al.* in rosettes. **f)** Summary table of crossed data obtained by mass spectrometry for the rosettes with the data from Ezzoukhry and *al.* paper.

**Supp Figure 3:**
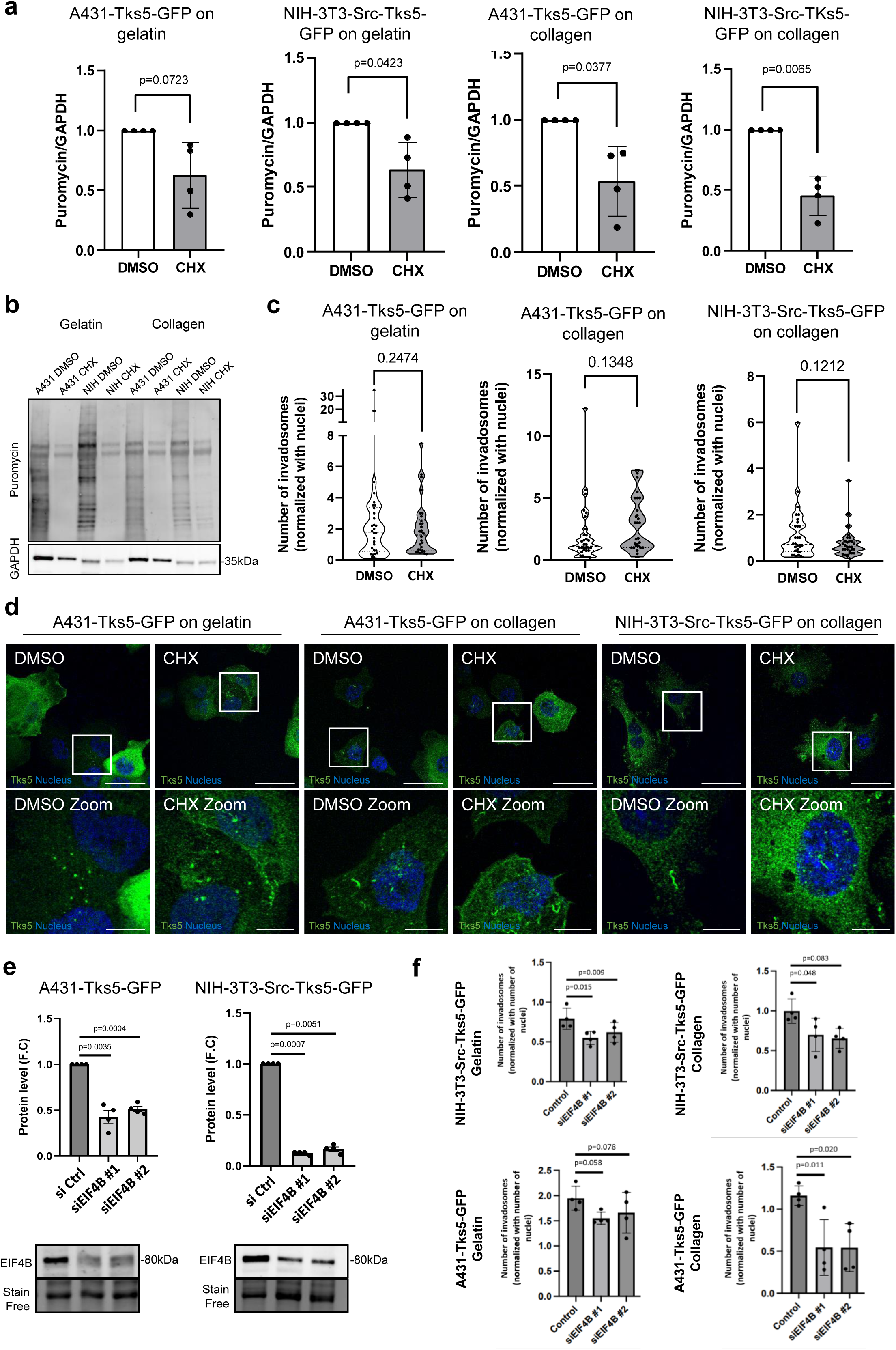
Translation is not involved on invadosome formation. **a)** Quantification of the effect of cycloheximide treatment on A431 and NIH-3T3-Src cells by using puromycin quantification. Values represent the mean +/- SEM of n=4 independent experiments and were analyzed using student t-test. **b)** Relative western blot analysis of puromycin expression for cells treated (CHX) or not (DMSO) with cycloheximide in A431 and NIH-3T3-Src cells seeded on gelatin or collagen. **c)** Quantification of the numbers of invadosomes per cell on gelatin and collagen treated (CHX) or not (DMSO) with cycloheximide in A431-Tks5-GFP and NIH3T3-Src-Tks5-GFP cells. Values represent the mean +/- SEM of n=3 independent experiments (10 images per condition and per replicate) and were analyzed using student t-test. **d)** Representative images of invadosome formation in A431-Tks5-GFP1 and NIH3T3-Src-Tks5-GFP cells seeded on gelatin or collagen. Tks5 is stained in green and nuclei in blue. Scale bar: 40µm, zoom: 10µm. **e)** Western Blot analysis of endogenous EIF4B expression in A431 and NIH-3T3-Src cells transfected using control or EIF4B-targeting siRNA and associated quantification. Stain free was used as the loading control. Values represent the mean of n=4 independent experiments. **f)** Quantification of the numbers of invadosomes per cell on gelatin and collagen silencing (siEIF4B) or not (DMSO) for EIF4B in A431-Tks5-GFP and NIH3T3-Src-Tks5-GFP cells. Values represent the mean +/- SEM of n=4 independent experiments (10 images per condition and per replicate) and were analyzed using student t-test.

**Video 1:**

Time-lapse video microscopy of lifeact-mRuby (red)-expressing NIH-3T3-Src cells transfected with KDEL-GFP (white).

**Video 2:**

Time-lapse video microscopy of lifeact-mRuby (red)-expressing NIH-3T3-Src cells transfected with KDEL-GFP (white).

**Video 3:**

Time-lapse video microscopy of lifeact-mRuby (red)-expressing NIH-3T3-Src cells transfected with KDEL-GFP (white).

**Video 4:**

Time-lapse of Lifeact-mRuby and KDEL-GFP signals recorded in NIH-3T3-Src cells treated with cycloheximide (CHX; 35μM).

**Supplemental table 1:**
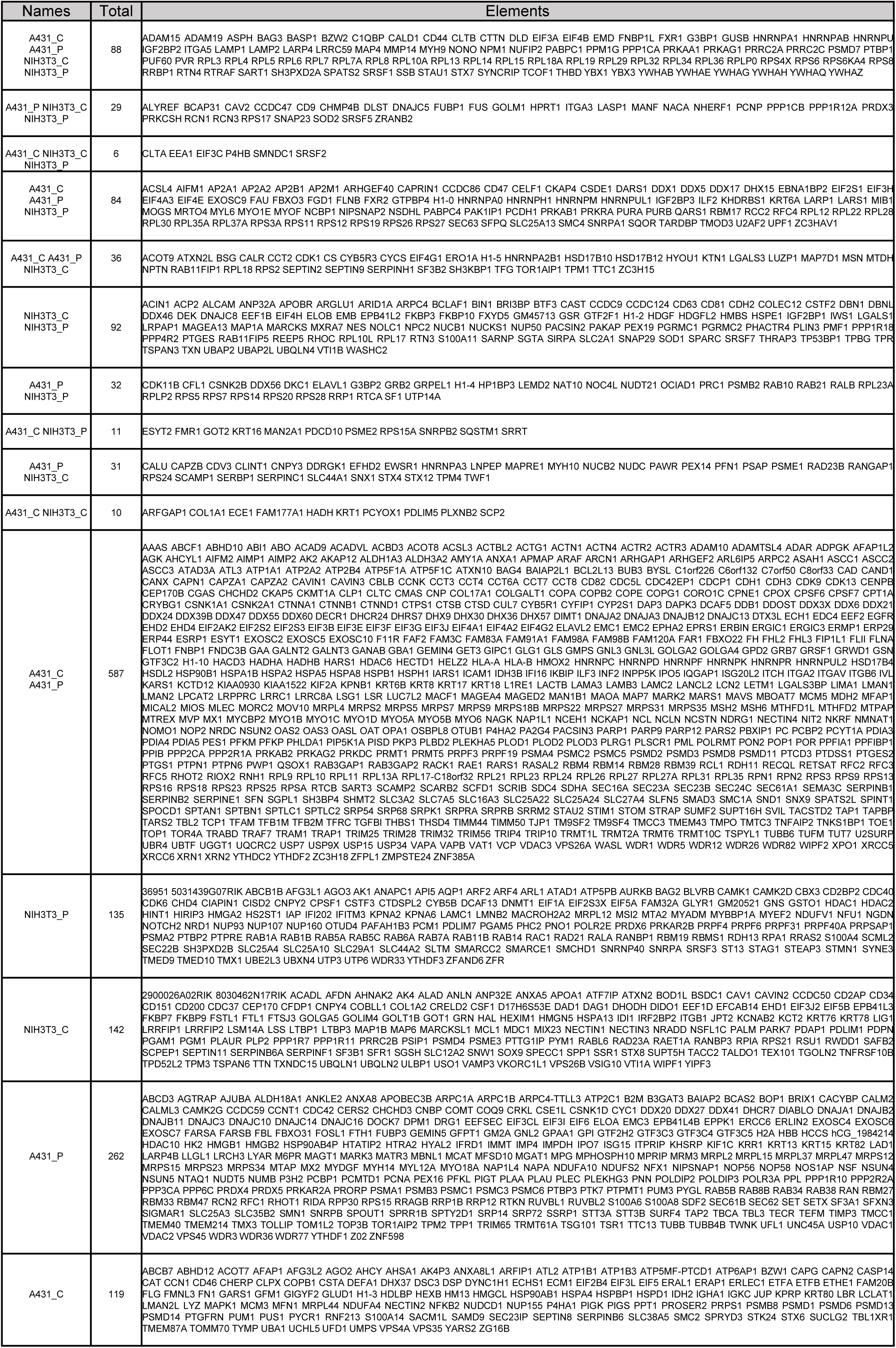
Summary table of proteins present in invadosomes identified by mass spectrometry analysis of Tks5 interactome

**Supplemental table 2:**
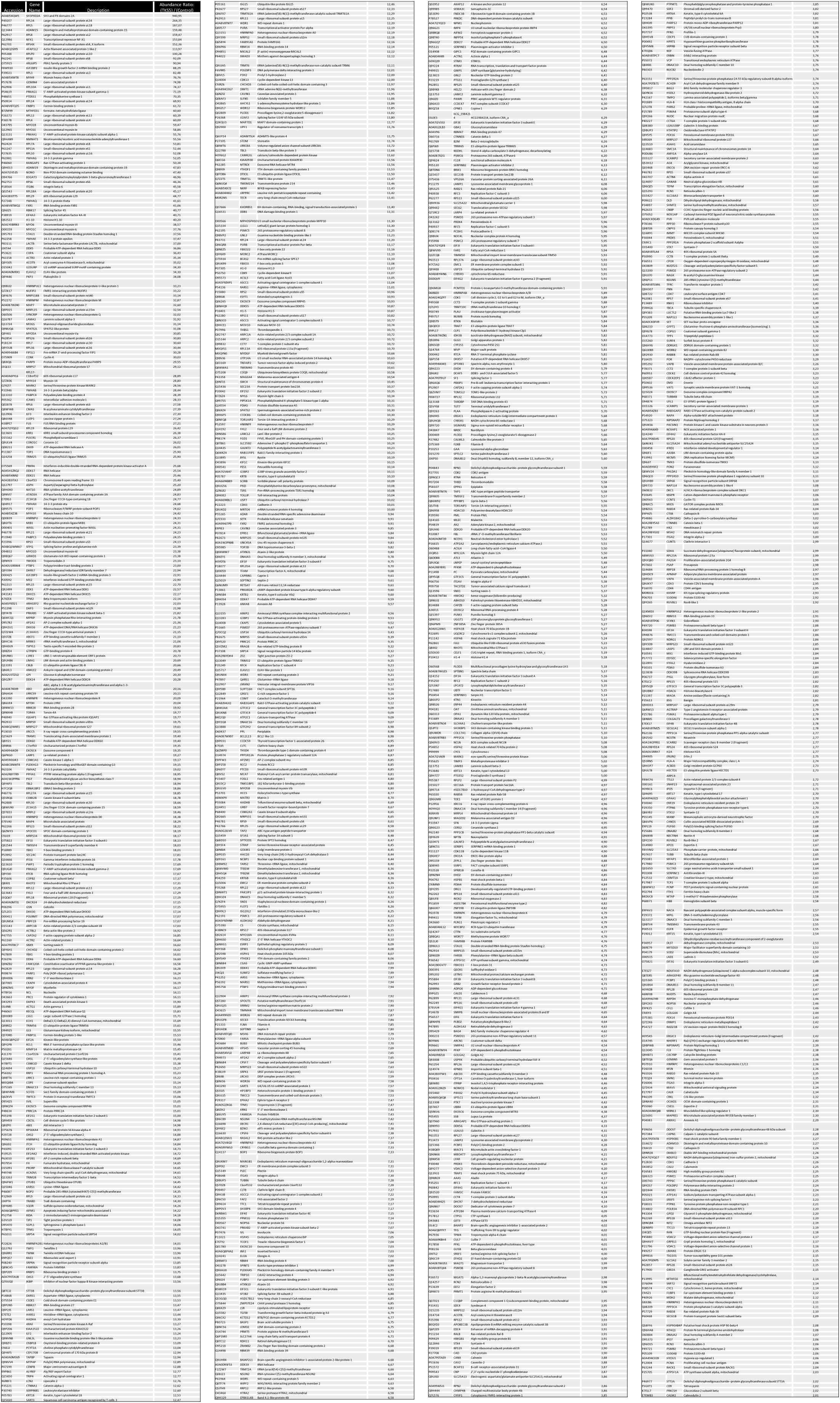
Summary table of proteins present in A431-Tks5-GFP cells seeded on plastic identified by mass spectrometry analysis of Tks5 interactome

**Supplemental table 3:**
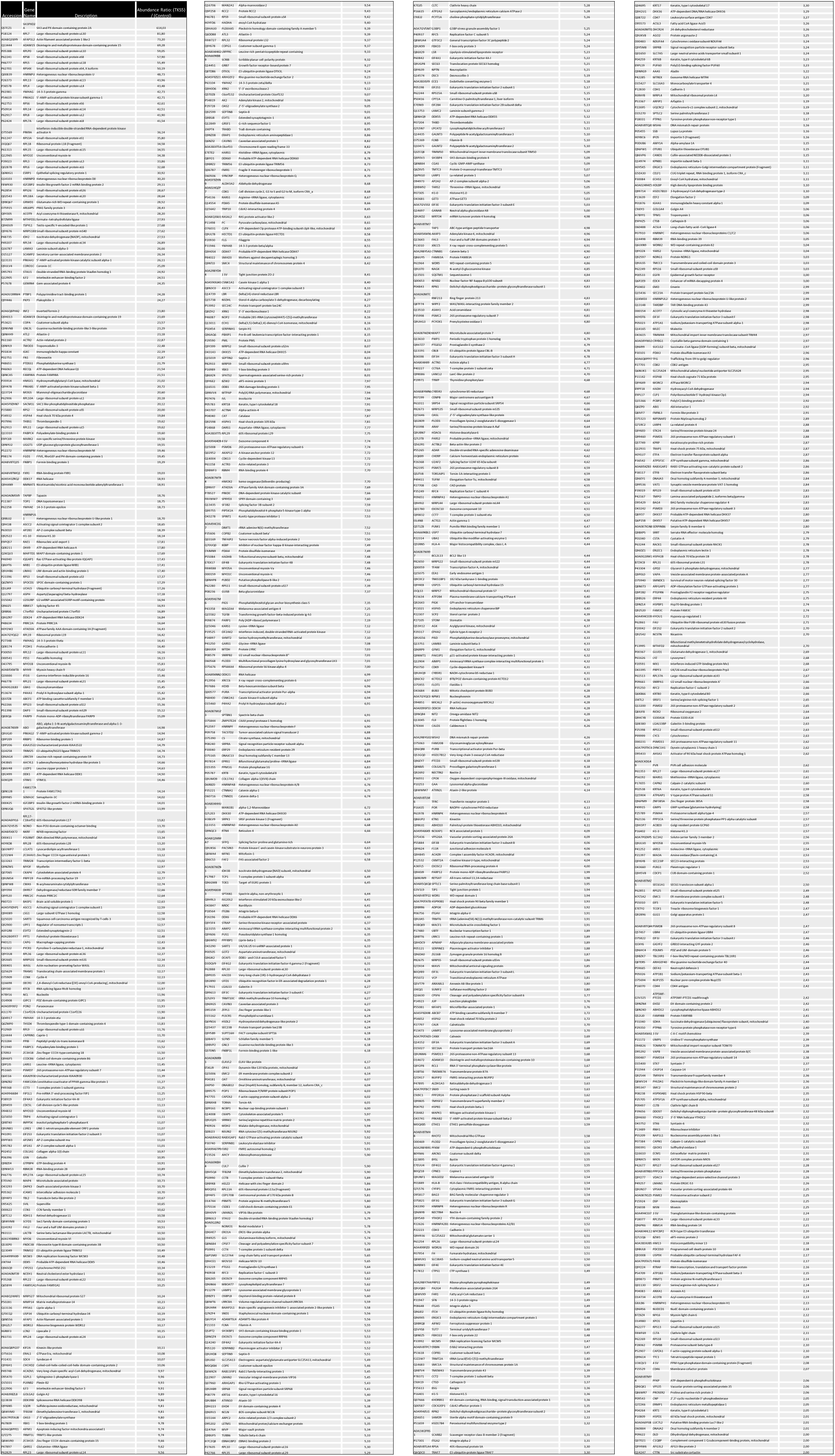
Summary table of proteins present in A431-Tks5-GFP cells seeded on collagen identified by mass spectrometry analysis of Tks5 interactome

**Supplemental table 4:**
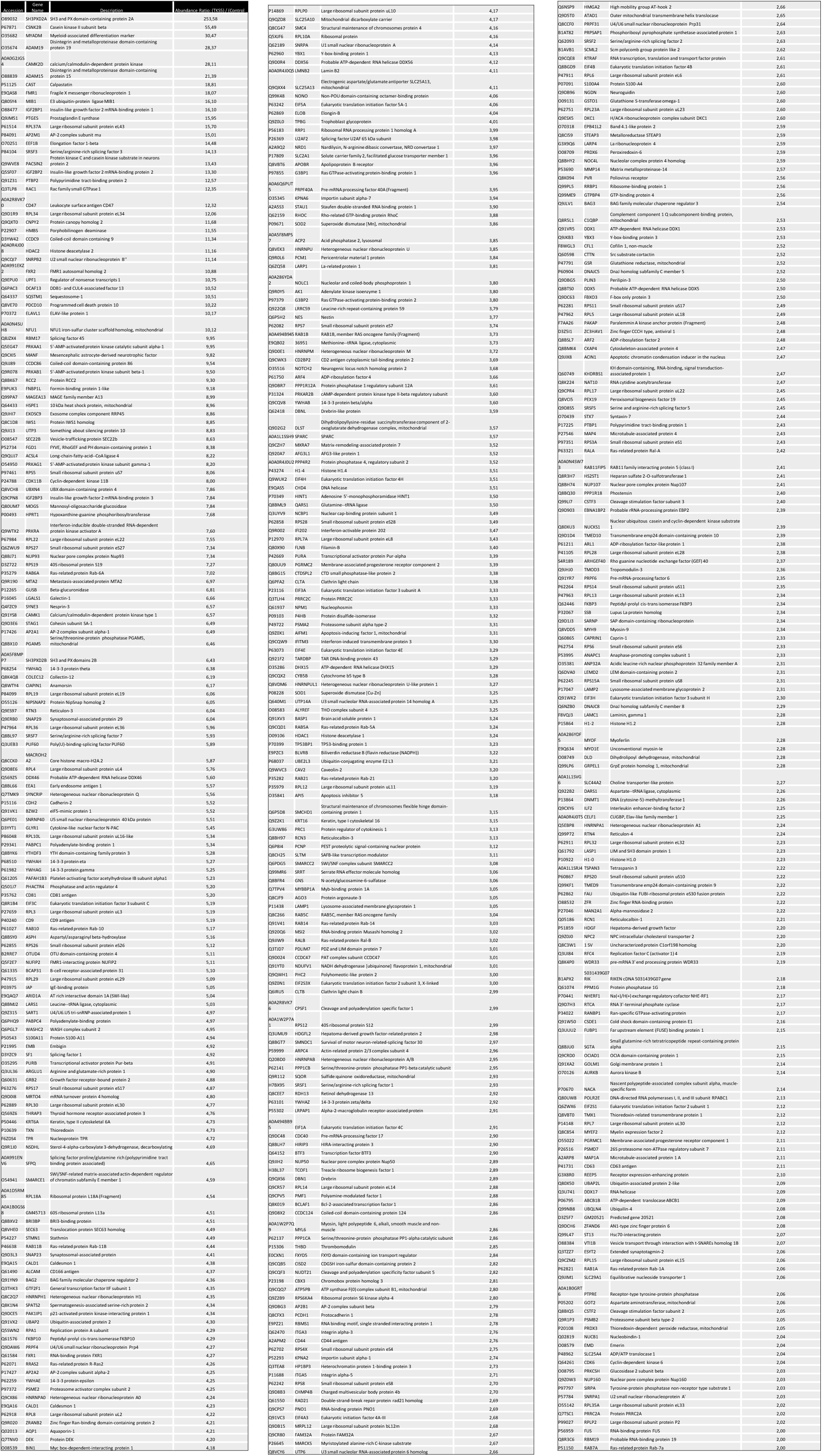
Summary table of proteins present in NIH-3T3-Src-Tks5-GFP cells seeded on plastic identified by mass spectrometry analysis of Tks5 interactome

**Supplemental table 5:**
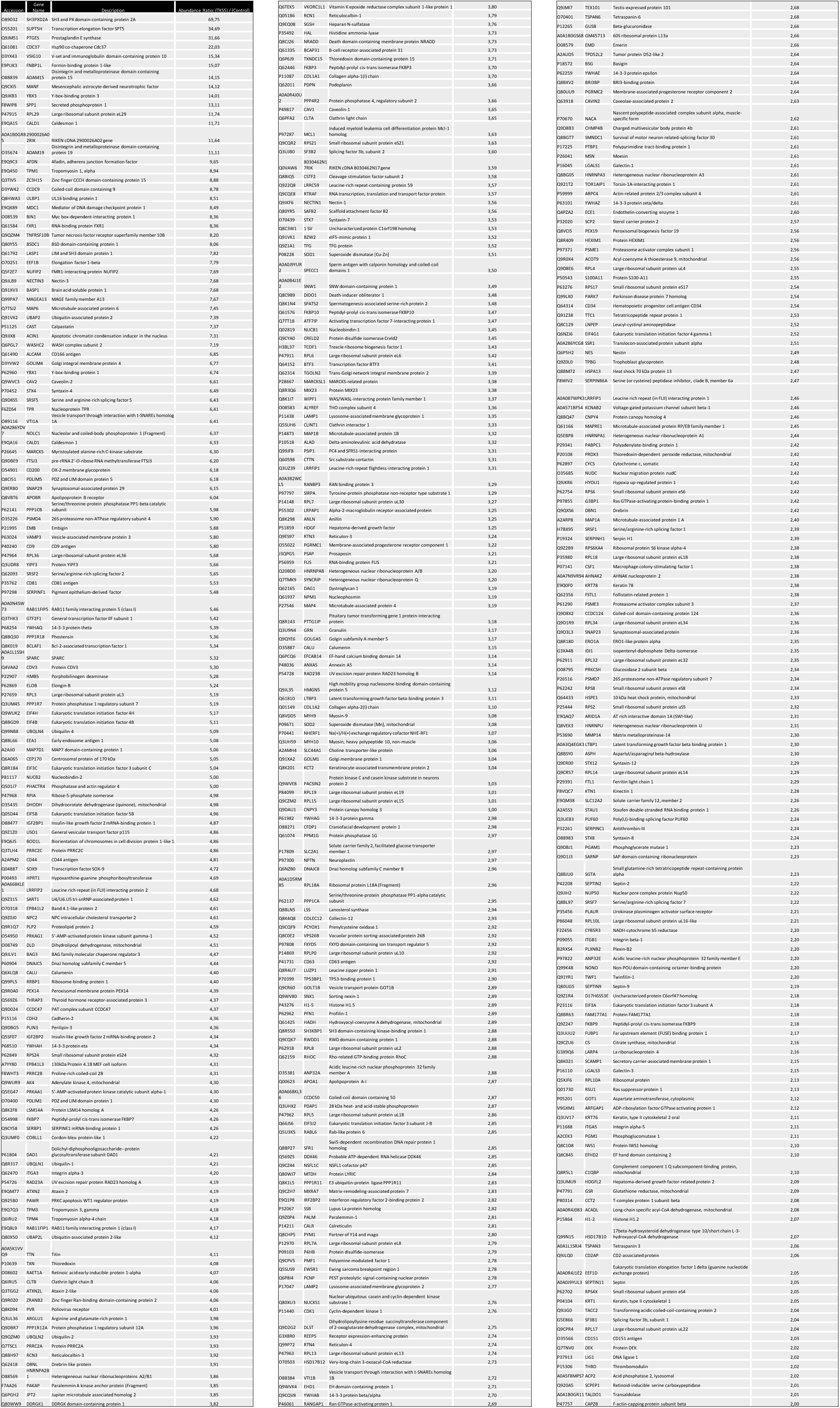
Summary table of proteins present in NIH-3T3-Src-Tks5-GFP cells seeded on collagen identified by mass spectrometry analysis of Tks5 interactome

## Notes

### Competing Interest Statement

The authors have declared no competing interest.

### Summary of Updates

The text has been edited for clarity. Discussion has been improved Experiments have been added (Figure 6b and Supp Fig 3g)

